# TEsorter: lineage-level classification of transposable elements using conserved protein domains

**DOI:** 10.1101/800177

**Authors:** Ren-Gang Zhang, Zhao-Xuan Wang, Shujun Ou, Guang-Yuan Li

**Affiliations:** Department of Bioinformatics, Ori (Shandong) Gene Science and Technology Co., Ltd., Weifang 261322, China; Shijiazhuang People’s Medical College, Shijiazhuang 050091, China; Department of Ecology, Evolution, and Organismal Biology (EEOB), Iowa State University, Ames, IA, 50010, USA

## Abstract

**Summary:** Transposable elements (TEs) constitute an import part in eukaryotic genomes, but their classification, especially in the lineage or clade level, is still challenging. For this purpose, we propose TEsorter, which is based on conserved protein domains of TEs. It is easy-to-use, fast with multiprocessing, sensitive and precise to classify TEs especially LTR retrotransposons (LTR-RTs). Its results can also directly reflect phylogenetic relationships and diversities of the classified LTR-RTs.

**Availability:** The code in Python is freely available at https://github.com/zhangrengang/TEsorter.

## 1 Introduction

Transposable elements (TEs) constitute the largest portion of most eukaryotic genomes, among which long terminal repeat retrotransposons (LTR-RTs) are predominant in plant genomes. Various tools have been developed for identification and classification of TEs or LTR-RTs, such as RepeatModeler (http://www.repeatmasker.org/RepeatModeler/), REPET (Quesneville, *et al*., 2005) and LTR_retriever (Ou and Jiang, 2017). To our knowledge, most of them can only classify TEs into the superfamily level, leaving the gap for revealing phylogenetic relationships between TEs, especially the LTR-RT *Copia* and *Gyspy* superfamilies. Previous studies (Llorens, *et al*., 2009; Neumann, *et al*., 2019; Wicker and Keller, 2007) have proposed classifications of LTR-RTs on lineage or clade levels. Particularly, Neumann *et al*. (2019) classified the *Copia* superfamily into *Ale, Alesia, Angela, Bianca, Bryco, Lyco, Gymco* I–IV, *Ikeros, Ivana, Osser, SIRE, TAR* and *Tork* lineages and the *Gypsy* superfamily into *CRM, Chlamyvir, Galadriel, Tcn1, Reina, Tekay, Athila, Tat* I–III, *Ogre, Retand, Phygy* and *Selgy* clades. These studies provide protein domain databases for lineage/clade-level LTR-RT classifications and moreover, the update of REXdb by Neumann *et al*. (2019) also provides classifications for other TEs, such as long interspersed nuclear repeats (LINEs), terminal inverted repeats (TIRs) and Helitrons. Here we take the opportunity to develop an automated, easy-to-use classifier, named TEsorter, to classify LTR-RTs as well as other TEs into detailed lineages/clades that reflect their phylogenetic relationships and diversities.

## 2 Methods

The TEsorter classifier was implemented using hidden Markov model (HMM) profiles obtained from protein domain databases GyDB (Llorens, *et al*., 2011) and REXdb (Neumann, et al., 2019). For REXdb, the viridiplantae v3.0 and metazoa v3 protein sequences were downloaded. Subsequently, multiple sequence alignments were performed by lineage and domain using MAFFT (Standley and Katoh, 2013) and HMM profiles were generated with HMMPress (Eddy, 1998).

Input DNA sequences were translated in all six frames and the translated sequences were searched against one of the two databases using HMMScan (Eddy, 1998). Hits with coverage < 20% or E-value > 1e-3 were discarded. For each domain of one sequence, only the best hit with the highest score was reserved. The classifications of TE superfamilies (e.g. LTR/*Copia*, LTR/*Gyspy*) and clades (e.g. *Reina* and *CRM* of *Gypsy*) were based on hits directly. For *Copia* and *Gyspy* superfamilies, complete elements were identified based on the presence and order of conserved domains including capsid protein (GAG), aspartic proteinase (AP), integrase (INT), reverse transcriptase (RT) and RNase H (RH) as described in Wicker *et al*. (2007). The identified domain sequences were extracted for further phylogenetic analyses.

To improve the classification sensitivity, a two-pass strategy was made available. The unclassified TE sequences were searched against the HMM-classified sequences using BLAST (Altschul, *et al*., 1990) and then classified with the 80–80–80 rule (Wicker, *et al*., 2007). This was based on the sequence-level similarity between autonomous and non-autonomous TEs, in which mutations like frameshifts and domain losses prevent their identification using HMMs. To comply with alignment uncertainties, this step only classified sequences at the superfamily level.

## 3 Results and Discussion

To benchmark the classification performance of TEsorter, we selected three non-redundant curated TE libraries from rice (Ou and Jiang, 2017), maize (Schnable, *et al*., 2009) and fruit fly (from Repbase v20.03, Bao, *et al*., 2015) and compared with four TE classifiers, including the RepeatClassifier module of RepeatModeler (http://www.repeatmasker.org/RepeatModeler/), the PASTEC module (Hoede, *et al*., 2014) of REPET, the annotate_TE module of LTR_retriever (Ou and Jiang, 2017) and the online-only LTRclassifier (Monat, *et al*., 2016). TEsorter with REXdb performed with the highest precision (0.94–1.0) in almost all the TE catalogs (Table 1, Supplementary Table S1). The sensitivity of TEsorter with REXdb was sub-optimal (0.79–0.93) in classifications of the LTR-RT *Copia* and *Gyspy* superfamilies in plants (Table 1). By searching against the Pfam database (Punta, *et al*., 2012), the unclassified LTR-RTs were confirmed to have lost their main protein domains. Some of these elements can be classified by using similarity to known elements. For this purpose, we implemented the two-pass strategy in TEsorter. However, due to the divergence of TE sequences, the homology-based approach only improved the sensitivity marginally (data not shown). As a result, a lower sensitivity was expected due to the rich of non-autonomous elements, including TIRs and Helitrons (Supplementary Table S1). In contrast, for autonomous TIR and Helitron elements, TEsorter performed much higher sensitivity (0.84–0.89) (Supplementary Table S1). TEsorter performed better with REXdb than with GyDB in plants (Table 1) due to the systematic collection of plant LTR-RTs by Neumann *et al*. (2019). Both databases showed low sensitivity (~0.5) in fly LTR-RTs classification (Supplementary Table S1), which might be a limitation of the domain-based approach on consensus sequences, as discussed by Monat, *et al*. (2016).

**Table 1.**
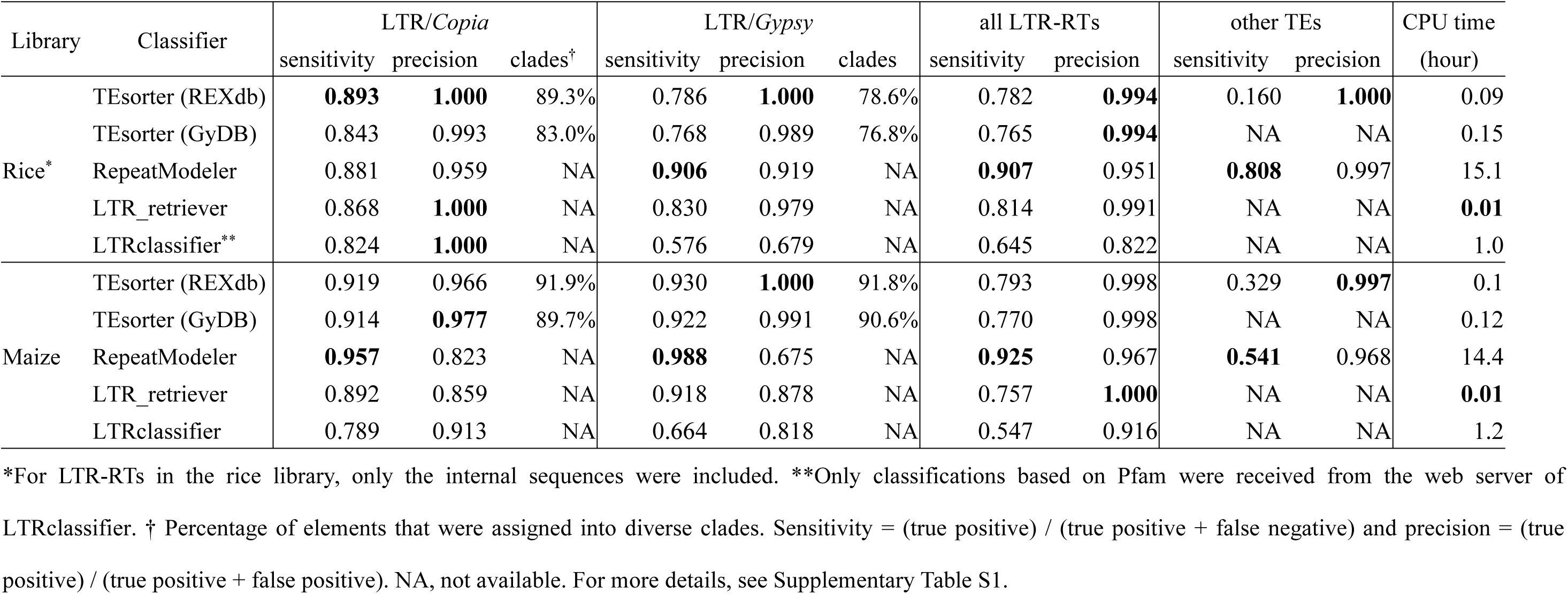
Comparison of the performance of difference classifiers.

| Library | Classifier | LTR/*Copia* |  |  | LTR/*Gypsy* |  |  | all LTR-RTs |  | other TEs |  | CPU time<br>(hour) |
| --- | --- | --- | --- | --- | --- | --- | --- | --- | --- | --- | --- | --- |
|  |  | sensitivity | precision | clades<sup>†</sup> | sensitivity | precision | clades | sensitivity | precision | sensitivity | precision |  |
| Rice* | TEsorter (REXdb) | **0.893** | **1.000** | 89.3% | 0.786 | **1.000** | 78.6% | 0.782 | **0.994** | 0.160 | **1.000** | 0.09 |
|  | TEsorter (GyDB) | 0.843 | 0.993 | 83.0% | 0.768 | 0.989 | 76.8% | 0.765 | **0.994** | NA | NA | 0.15 |
|  | RepeatModeler | 0.881 | 0.959 | NA | **0.906** | 0.919 | NA | **0.907** | 0.951 | **0.808** | 0.997 | 15.1 |
|  | LTR_retriever | 0.868 | **1.000** | NA | 0.830 | 0.979 | NA | 0.814 | 0.991 | NA | NA | **0.01** |
|  | LTRclassifier** | 0.824 | **1.000** | NA | 0.576 | 0.679 | NA | 0.645 | 0.822 | NA | NA | 1.0 |
| Maize | TEsorter (REXdb) | 0.919 | 0.966 | 91.9% | 0.930 | **1.000** | 91.8% | 0.793 | 0.998 | 0.329 | **0.997** | 0.1 |
|  | TEsorter (GyDB) | 0.914 | **0.977** | 89.7% | 0.922 | 0.991 | 90.6% | 0.770 | 0.998 | NA | NA | 0.12 |
|  | RepeatModeler | **0.957** | 0.823 | NA | **0.988** | 0.675 | NA | **0.925** | 0.967 | **0.541** | 0.968 | 14.4 |
|  | LTR_retriever | 0.892 | 0.859 | NA | 0.918 | 0.878 | NA | 0.757 | **1.000** | NA | NA | **0.01** |
|  | LTRclassifier | 0.789 | 0.913 | NA | 0.664 | 0.818 | NA | 0.547 | 0.916 | NA | NA | 1.2 |

\*For LTR-RTs in the rice library, only the internal sequences were included. \*\*Only classifications based on Pfam were received from the web server of LTRclassifier. † Percentage of elements that were assigned into diverse clades. Sensitivity = (true positive) / (true positive + false negative) and precision = (true positive) / (true positive + false positive). NA, not available. For more details, see Supplementary Table S1.

RepeatClassifier had the best sensitivity in most cases (Table 1, Supplementary Table S1), which was benefited from Repbase that has collected TE sequences from the three species we benchmarked. PASTEC in the REPET pipeline also uses Repbase for classification. However, it only provided confident classifications at the order level (Supplementary Table S1). LTRclassifier and LTR_retriever used a set of selected Pfam domains for LTR-RT classifications. However, the selected Pfam domains aim for broad representation instead of clade-specific classification. TEsorter generally exhibited higher sensitivity and precision comparing to these two methods (Table 1, Supplementary Table S1).

TEsorter assigned 76–92% of LTR *Copia* or *Gypsy* elements into diverse clades in plants (Table 1). We performed phylogenetic analyses to evaluate the precision of these clade-level assignments. Briefly, protein domain sequences were extracted using TEsorter and aligned with MAFFT (Standley and Katoh, 2013), and the phylogenetic trees were reconstructed using IQ-TREE (Nguyen, *et al*., 2015). Using RT domains as an example, the clade-level classification of TEsorter was highly consistent (99.06%) with the phylogeny (Supplementary Fig. S1a) and also consistent with the previous report (Neumann, *et al*., 2019). Similar high consistencies were observed on other domains’ classification (Supplementary Fig. S1b–d). These results revealed high-confidence classifications at the clade level by TEsorter.

The TEsorter package was implemented in Python and was accelerated using multiprocessing (Table 1).

---

**Supplementary Fig. S1. Consistency between classifications of TEs and phylogenetic relationships based on RT (a), RH (b), INT (c) and concatenated RT**–**RH**–**INT (d) domains in rice.** Conflicts were highlighted by black circle nodes. The tree was un-rooted. Branches were colored based on TEsorter classifications.

**Supplementary Table S1. Performances with different TE catalogs.**

## Acknowledgments

We thank Dr. Neumann Pavel for the notification of the release of REXdb and Dr. Jia-Hui Chen for suggestions to analyze RT domains of LTR-RTs and LINEs.

