## Supplementary figures and images for "TEsorter: lineage-level classification of transposable elements using conserved protein domains"

### Supplementary Fig. S1

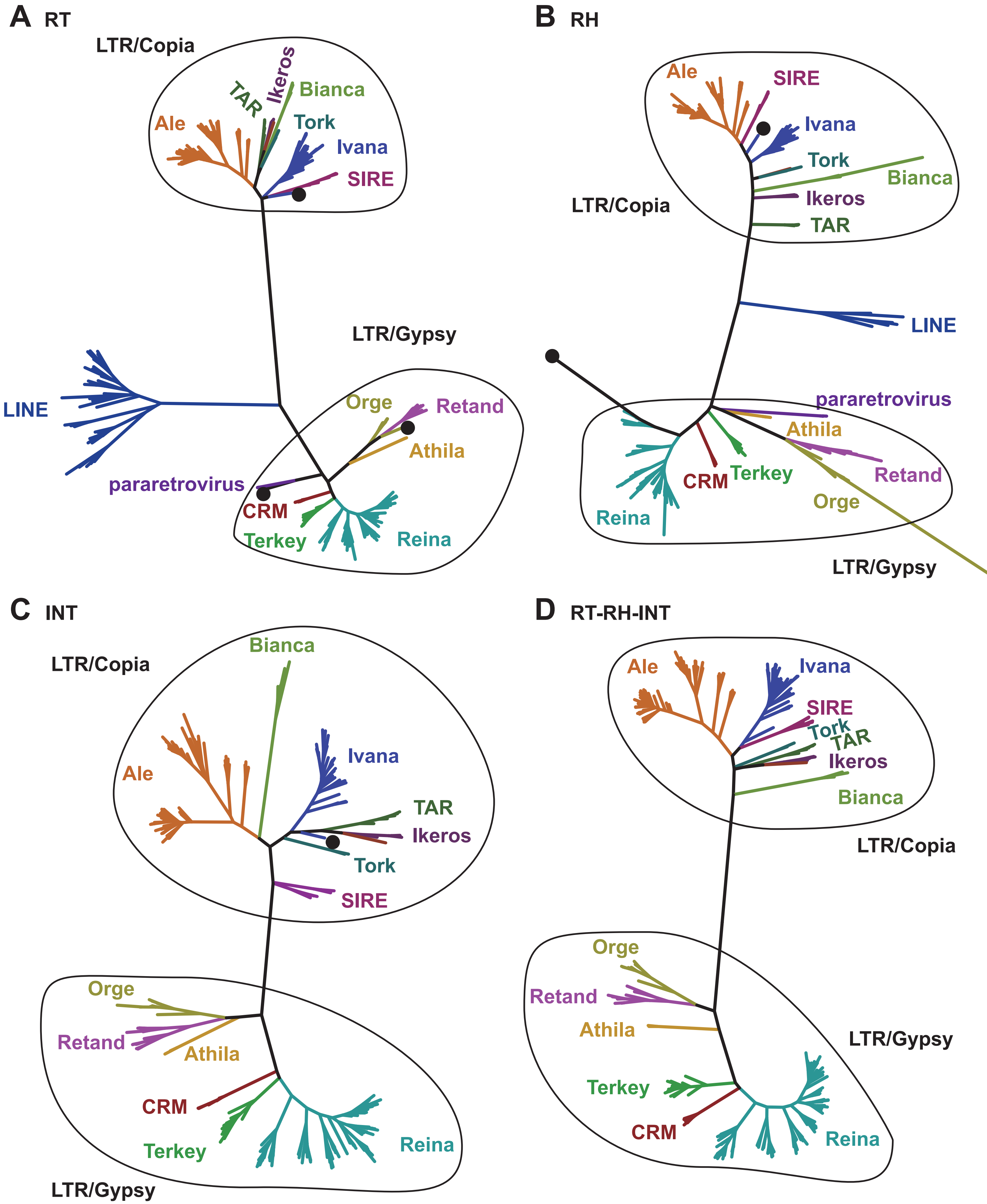
